# Validation of a Commercial Radioimmunoassay for Measurement of Human Peripheral Oxytocin

**DOI:** 10.1101/2023.03.22.533823

**Authors:** Keenan Gerred, Amita Kapoor

## Abstract

Oxytocin (OT) is a peptide hormone synthesized in the hypothalamus and released into systemic circulation or other areas of the brain. It has a variety of physiological roles such as stimulation of uterine contractions, involvement in social behaviors, and regulation of mood. Its small size and low levels within biological matrices make it challenging to accurately measure. The goal of this study was to demonstrate the specificity of the antibody, sensitivity, and reproducibility of the Phoenix Pharmaceuticals (PP) OT radioimmunoassay (RIA) for use in human urine, serum, and saliva. Specificity of the antibody was assessed by high pressure liquid chromatography with ultraviolet (HPLC-UV) separation and assay of the fractions. Immunoreactivity was evaluated using the percent OT bound, and the fraction retention times were compared to the retention time of an intact OT standard to determine which fractions contained OT in the extracted samples. Reproducibility was assessed by running replicates of pools of each biomatrix over several assays. Sensitivity was assessed by repeated measurement of physiologically relevant low-concentration specimens. In all tested specimens the greatest reactivity in assay corresponded to the same fraction(s) as the OT standard. Only minimal reactivity was found in the other fractions, suggesting that in an unfractionated sample the antibody reacts mostly with intact OT. Reproducibility was acceptable for all specimens and the coefficient of variation (CV) ranged from 3.72-8.04% and 5.89-12.8%, for intra and inter-assay, respectively. The limits of quantitation (LOQ) were sufficient for measurement of normal values in urine (0.643 & 1.43 pg/mL), serum (1.90 pg/mL), and saliva pools (0.485 & 4.42 pg/mL). In conclusion, the PP OT RIA is specific and sensitive enough for reproducible measurement of intact OT in human peripheral biological matrices.

**HIGHLIGHTS:**

- The oxytocin radioimmunoassay kit from Phoenix Pharmaceuticals is specific for intact oxytocin measured in human urine, serum, and saliva.
- Basal levels of oxytocin are low in peripheral biomatrices, and the sensitivity of the assay is sufficient for reproducible measurement.

## 1. INTRODUCTION

Oxytocin (OT) is a nine-amino acid peptide hormone that is primarily synthesized in the magnocellular neurons of the paraventricular and supraoptic nuclei of the hypothalamus ^1^. OT is released into systemic circulation via the posterior pituitary and into the central nervous system via widespread OT-ergic pathways ^2^. OT has well-known physiological functions during labor and lactation, and it acts as a neuromodulator and plays a role in social cognition, executive functions, maternal behavior, brain development, pain perception, bonding, and others ^3–6^. Given these roles, direct measurement of central OT levels is desirable, however; collection of cerebrospinal fluid (CSF) to directly measure these levels is rarely feasible. Peripheral measures of OT have thus traditionally been used as a surrogate including blood (serum, plasma) as well as alternative biological matrices such as saliva and urine. These peripheral measures have been used in a wide range of studies which include obesity and metabolism ^7^, neuropsychiatric disorders ^8^, and social behaviors ^9^.

There is much controversy surrounding the measurement of OT (e.g., ^10–12^). First, sample preparation techniques are not consistent across studies and have led to wildly different OT concentrations. For example, assaying unextracted samples resulted in values that were 10-1000 times higher compared to extracted samples ^13^. Presumably this was due to cross-reactivity with interfering compounds and/or matrix effects since the samples were not purified. A second issue with OT measurement is that commercially available assays yield different results, even when interfering substances are removed and the same assays are utilized across studies^23^. This could be due to the specificity of the antibodies, issues with sensitivity & matrix effects, recognition of differing epitopes, and/or the conformational state of the OT molecule ^11,14^. The small size & simple structure of OT also makes it difficult to produce antibodies with sufficient reactivity & specificity^23^. A third issue is the low levels of endogenous OT in peripheral matrices with the normal levels ranging from 0-10 pg/mL(1-3pg/mL in blood), which can lead to mostly non-detectable values by any currently-available measurement technique ^12,15^. OT is promptly cleared out of the blood upon peripheral release: it has a short half-life of 1-5 minutes and is rapidly degraded by peptidases in the kidney and liver ^16^. Results are inconsistent regarding the levels of intact OT that can be found in urine, as some studies suggest that urine does not contain intact OT, but others suggest that there is approximately 1% clearance of OT into the urine ^16,17^. The half-life of endogenous OT in saliva is unknown but is estimated to be close to that in blood ^18^. Further adding to the controversy, the biological relevance of peripheral OT and its relationship to central OT measures are unclear, however; this is not the focus of this paper ^19^.

Traditionally, OT was measured by radioimmunoassays (RIA) for which full in-house method validations were conducted ^10^. However, when commercially available enzyme immunoassays (EIA) for measurement of OT entered the market, they were rapidly adopted as they were more accessible, faster, and did not use radioactivity. These assays did not always undergo the fit-for-purpose validations that would make their results relevant to performance of the assays in studies with normal levels in peripheral biological matrices. This has led to questionable performance of some often-utilized kits when measuring endogenous OT peripherally ^10^. Many validations have used administration of intranasal OT to demonstrate that the assays detect changes in OT concentrations, but this is not appropriate for endogenous peripheral measurement since the levels produced are not within the endogenous concentration range, and cross-reactivity of antibodies with any absorbed intranasal OT confounds the results. More recently, assays have been developed that measure OT using liquid chromatography-mass spectrometry (LC-MS/MS) which is considered the gold-standard for measurement as the detection is highly specific ^15,20^. The LC-MS/MS methods tend to have concentration ranges like the original OT RIAs.

The goal of this study was to determine if there was a commercially available immunoassay kit to specifically and reproducibility measure OT as an alternative to LC-MS/MS methodology. To this end, we identified an RIA by Phoenix Pharmaceuticals and preliminary evaluation of this kit demonstrated values that were physiologically relevant, reproducible, and in line with those obtained by LC-MS/MS and older RIA methodology. As researchers are generally interested in peripheral levels of OT, the validation focused on urine, serum, and saliva.

## 2. METHODS

### 2.1 Preparation of pooled sample material

Human donor material was obtained for each biological matrix for the validation experiments. Urine was collected in-house from volunteers and immediately acidified with 10µL/mL of concentrated phosphoric acid. Pooled human serum was obtained from Innovative Research (Novi, Michigan) and 0.6 TIU/mL of recombinant aprotinin was added (Sigma; Burlington, MA). For saliva, two large-volume samples from different donors were purchased from Innovative Research and combined. All the pools were appropriately aliquoted and stored at -80°C before use.

### 2.2 Solid phase extraction

Solid-phase extraction (SPE) was carried out prior to assay. Acetonitrile (ACN), methanol (MeOH), and trifluoroacetic acid (TFA) were purchased from Fisher (Fisher; Waltham, MA). Ultrafiltered water (UF H_2_O) was used for all solutions and was produced in-house using an US Filter PureLab PLUS UV/UF filtration system. SPE cartridges for all extractions were Waters Sep-Pak Vac 3cc 200mg C18 (Waters; Milford, MA).

#### 2.2.1 Urine

The acidified aliquots were thawed on ice, vortexed, and centrifuged (10 min, 1,000xG, 4°C). The SPE columns were conditioned with 3mL of 100% MeOH followed by 3mL of UF H_2_O, and then the sample (3mL) was loaded. Pressure was briefly applied before the samples were allowed to load by gravity for 10 minutes before completion with pressure. The cartridges were then washed with 3 × 3mL 10% ACN in 1% TFA, followed by elution with a 3mL aliquot of 80% ACN for 10 minutes by gravity before completion with pressure. The solvent was dried in a centrifugal concentrator and stored at -20°C until analysis.

#### 2.2.2 Serum and saliva

Aliquots (1mL) were thawed on ice before the addition of 1mL of chilled 1% TFA, vortexed, and then centrifuged (20 mins, 17,000xG, 4°C). The samples were then loaded onto SPE cartridges conditioned with 1mL of 60% ACN in 1% TFA followed by 3×3mL 1% TFA. Pressure was briefly applied before the samples were allowed to load by gravity for 10 minutes and completed with pressure. The cartridges were then washed with 2×3mL 1% TFA and OT was eluted with a 3mL aliquot of 60% ACN in 1% TFA which was allowed to flow by gravity for 10 minutes before completion of the elution with pressure. The solvent was then dried in a centrifugal concentrator and stored at -20°C until HPLC analysis.

### 2.2 HPLC-UV

HPLC separation was performed on a Thermo-Scientific Vanquish system with a diode array UV detector. The mobile phase was isocratic 20% ACN in 0.08M phosphate buffer pH 5.5 for 6 minutes, followed by a wash with 95% ACN before re-equilibration with mobile phase. The column was a Phenomenex Gemini 150 × 4.6mm C18 (00F-4435-E0). Standards for OT and Arg-8 vasopressin (AVP) were purchased from Sigma. AVP was included for determination of specificity as it is very similar to OT and is a common cross-reactant with the antibodies in some commercially available immunoassays for OT.

For determination of the elution positions of OT and AVP, the compounds were reconstituted at 1mg/mL in HPLC-grade water (Fisher W5-4) and stored in aliquots at -80°C. On the day of analysis, the standards were diluted to 1µg/mL with 20% ACN and then 20µL of each was injected and the resultant chromatograms were evaluated to identify the OT and AVP standard peaks.

For fractionation the extracted samples from each biomatrix were reconstituted in 30µL of 20% ACN before injection of 20µL on the column. 5µL of an OT reference standard solution was also added to a spiked urine sample prior to injection. After injection, 0.5mL fractions were collected for 6 minutes using a Gilson FC-203B fraction collector for a total of 12 fractions. The fractions were then dried in a centrifugal concentrator and stored at -20°C until immunoassay. Those fractions that would contain AVP or OT were identified based on the retention time of the standards run that day.

### 2.3 Radioimmunoassay

The Phoenix Pharmaceuticals (Burlingame, CA) Oxytocin Radioimmunoassay (RIA) kit (catalog # RK-05-01) was used for immunoassay analysis. It is a competitive liquid-phase double-antibody RIA utilizing I-125 labeled tracer and was performed according to the kit protocol. The HPLC fractions were reconstituted in 250µL of assay buffer, and 100µL of each sample, standards prepared in assay buffer, and primary antibody were incubated in duplicate in polystyrene 12×75mm tubes overnight for 20-24 hours at 4°C.

On the second day of the assays the Stock Tracer was diluted in the assay buffer to a working counts-per-minute (CPM) of ∼10,000CPM using a 5-minute count time. 100µL of the Working Tracer was then added to all tubes before a second 20-24 hour incubation at 4°C.

On the third day of the assay 200µL of secondary antibody solution was added to the tubes before incubation at room temperature for 90 minutes, addition of 500µL assay buffer, and centrifugation before aspiration of the liquid. The CPM of the remaining pellet was counted on a Perkin-Elmer WIZARD2 gamma counter for 5 minutes. A % bound value was then obtained by dividing the mean NSB-corrected CPM of each standard and sample by the mean NSB-corrected CPM of the 100% binding tubes and multiplying by 100. The values of the standards were plotted against the % bound to obtain a standard curve using a 4-parameter logistic fit, and the % bound values for the samples were interpolated to obtain the concentrations. Serum & saliva samples were concentrated by a factor of 4 and urine by a factor of 12 through extraction. Competitive assays measure the displacement of binding of conjugate from the antibody, therefore the more OT there is in the sample the less conjugate can bind, and therefore the % bound value produced by the assay is lower.

### 2.4 Data Analysis & Statistics

The % conjugate bound output by the assay was subtracted from 100 to obtain a theoretical % OT bound for use in analysis, and this as well as the OT levels in the fractions were evaluated to determine which had reacted in the immunoassay and graphed vs chromatographic time. Peaks were visually observed to determine if they aligned with the pure OT standard & the exact retention times were also compared. Sensitivity of the assay was assessed by diluting the pooled samples to determine the concentration where the CV was reliably under 15% and above 20% for the LOQ and LOD, respectively. Prism (9.5.0) and MyAssays Desktop Pro were used for analysis.

## 3. RESULTS

### 3.1 Specificity

Pooled samples were separated by HPLC-UV and the fractions collected. A standard mixture of AVP and OT was injected, and they eluted approximately in fractions 4 and 7-9, respectively (Figure 1A). The fractions from each biological matrix were measured for OT using the PP RIA and Figures 1B-D demonstrated that the largest amount of immunoreactivity corresponded to the fraction(s) that had the OT standard in all matrix types. A slight delay in elution timing was observed in the urine sample and confirmed by fractionating & running a spiked urine sample in assay.

**Figure 1:**
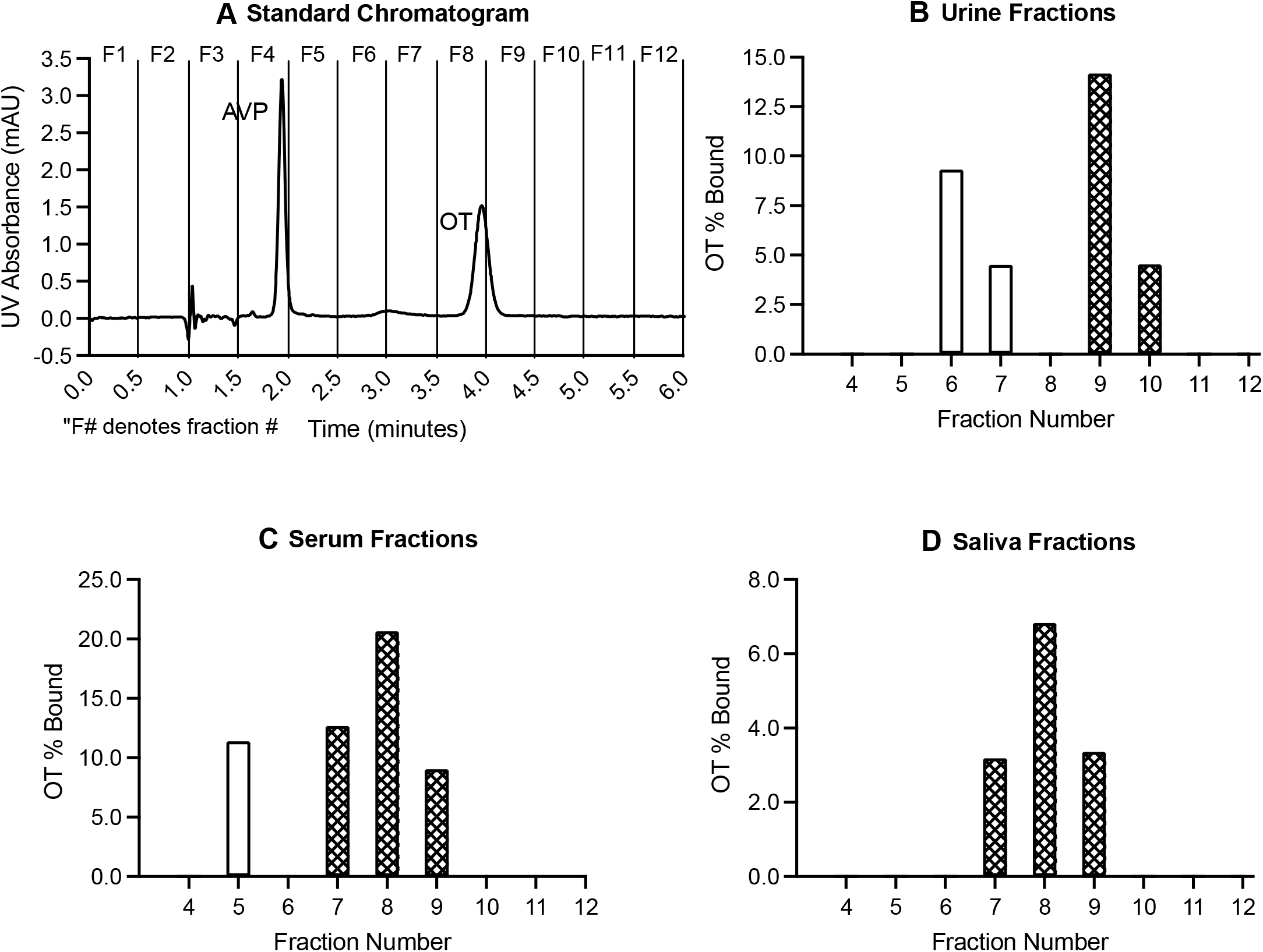
**1A)** Example chromatogram of standard injections of arginine vasopressin (AVP) and oxytocin (OT), the fraction numbers represented in the samples are labeled above the graph. **1B)** OT immunoreactivity of the fractions separated by HPLC-UV in urine, **1C)** serum, and **1D)** saliva. Patterned bars represent the elution position of the OT standard.

### 3.2 Reproducibility & Sensitivity

Reproducibility of the assay was demonstrated by running replicates (range: 11-40) of the pooled samples over 3-4 assays (Table 1). Intra-assay CV was under 10% and inter-assay CV was under 15% for all biological matrices at all concentration levels. Urine data is typically presented as pg/mg of creatinine to correct for urine dilution and those data were 0.753 pg/mg for the Urine 1 pool and 0.637 pg/mg for the Urine 2 pool. Sensitivity was calculated from repeated measurements of diluted pool samples created by reconstitution in differing amounts of assay buffer. The LOQ was at the point of the lowest calibrator and the LOD of the assay in the various biological matrices was slightly lower (Table 2).

**Table 1:** Mean OT concentration, intra and inter assay coefficients of variation (CV) of pooled serum, urine, and saliva samples.

| Biological Matrix | OT mean pg/mL | Intra-assay CV % (n assays) | Inter-assay CV % (n replicates) |
| --- | --- | --- | --- |
| Serum | 1.90 | 3.72 (3) | 5.89 (31) |
| Urine 1 | 1.43 | 6.71 (4) | 9.85 (40) |
| Urine 2 | 0.643 | 8.04 (4) | 11.63 (38) |
| Saliva 1 | 0.485 | 5.85 (4) | 12.8 (11) |
| Saliva 2 | 4.42 | 4.06 (4) | 7.89 (12) |

**Table 2:** Measured limit of quantitation (LOQ) and limit of detection (LOD) for each biological matrix corrected for volume.

| Biological Matrix | LOQ (pg/mL) | LOD (pg/mL) |
| --- | --- | --- |
| Urine | 0.104 | 0.049 |
| Serum | 0.313 | 0.207 |
| Saliva | 0.313 | 0.274 |

## 4. DISCUSSION

This study has shown that the PP OT RIA has sufficient specificity and reproducibility to use for reliable measurement of OT in serum, urine, and saliva ^21^. For all samples, the measured OT reactivity was contained within the fractions that contained the OT standard.

Reproducibility data showed that the CVs were within an acceptable range of less than 10% and 15%, for intra and inter assay, respectively; and as expected reproducibility was decreased as the concentration approached the LOQ. Furthermore, the % bound of the LODs ranged between 85-95% which is typical for competitive RIAs ^22^.

All the sample types also had reactivity in the assay with fractions that did not correspond to the OT standard, and this was especially prevalent in urine. There are a number of explanations for this finding. First, the nature of the competitive immunoassay will lead to increased interference of matrix and non-specific components if the analyte is not present.

Since the OT in the fractionated samples was isolated to certain fractions, it was not unexpected to observe higher background binding in the other fractions. Second, there was likely some matrix interaction as we saw higher levels of this background binding in urine samples compared to serum and saliva as it is more difficult to remove the interfering components found in urine by SPE. Some amount of interference remaining in urine is found regardless of extraction or assay methodology. All of these comparisons only included data from samples that were extracted as others have demonstrated that measurement of OT in non-extracted samples results in inflated values, presumably due to non-specific binding ^13,23^.

It is noteworthy that the concentration of OT in all biological matrices was low, so the samples needed to be concentrated through extraction to measure in the calibration range, a feature typical of most modern OT assay workflows. Early studies using RIAs demonstrated that basal levels of OT in plasma or serum had mean values that were approximately 1-3 pg/ml ^17,24^. Similarly, more recent LC-MS/MS data showed that levels ranged from below the level of detection (<1pg/ml) to 3 pg/ml ^15,25^. The mean level of OT in the current study using a serum pool was 1.90 pg/ml showing that this assay is consistent with other data, including gold standard mass spectrometry. These levels contrast with many EIA kits in which levels ranged from 4.82-7.53 pg/ml when the sample was extracted and suggests that these EIAs had less specificity.

There have only been a few studies that have measured saliva OT using methods other than EIA kits. As saliva as a biological matrix rose to popularity in the early 2000s, to our knowledge there are no studies that measured basal saliva OT using the original RIAs. Using LC-MS/MS, basal levels of human saliva were not-detectable ^15^. This was the case with the current study as we were not able to measure basal levels of OT in the commercial pool and needed to spike with OT standard to do the reproducibility experiments. Since spiked saliva was quantifiable at 0.485 pg/ml with acceptable reproducibility, it follows that basal saliva levels may be lower than that. Modern, ultra-sensitive assay techniques should be used for basal measurements in saliva (newer mass spectrometry, electrochemiluminescence, or Simoa).

Urine is an especially challenging biological matrix due the presence of metabolites & matrix components such as salts, which can decrease assay performance, even with extensive sample preparation techniques. Indeed, in this study we noted that the chromatograms of the urine matrix had more background and non-specific reactivity than the other matrices.

Nonetheless, using the PP OT RIA, we were consistently able to measure basal urine levels with acceptable reproducibility, suggesting that it is less susceptible to the interferences. Most other studies that have measured basal urine OT levels used EIA and found that levels ranged from 9.81-13.2 to even hundreds of pg/ml which is substantially higher than the current study and the expected levels in blood and CSF. Even using traditional RIAs the range was 2.5-34.068 pg/ml, suggesting non-specific binding with many immunoassay techniques ^13,17,26,12,15,20^, with the issues summarized in Tabak 2022. This makes consistent measurement of basal levels of OT in urine difficult, and caution should be used when measuring & comparing basal levels, even with the assay tested in this study.

In conclusion, we have shown that although there are numerous challenges to measuring peripheral OT due to the characteristics of the molecule and its low levels it can be reproducibly and reliably measured using the PP OT RIA. The kit is appropriate to use for quantification of peripheral OT in serum, urine, and saliva.

## 8. ACKNOWLEDGEMENTS

This work was supported by the Wisconsin National Primate Research Center P51 grant.

